# COSPLAY: An expandable toolbox for combinatorial and swift generation of expression plasmids in yeast

**DOI:** 10.1101/630277

**Authors:** Youlian Goulev, Audrey Matifas, Vincent Heyer, Bernardo Reina-San-Martin, Gilles Charvin

## Abstract

A large number of genetic studies in yeast rely on the use of expression vectors. To facilitate the experimental approach of these studies, several collections of expression vectors have been generated (YXplac, pRS series, etc.). Subsequently, these collections have been expanded by adding more diversity to many of the plasmid features, including new selection markers and new promoter sequences. However, the ever growing number of plasmid feature makes it unrealistic for research labs to maintain an up-to-date collection of plasmids.. Here, we developed the COSPLAY toolbox: a Golden Gate approach based on the scheme of a simple modular plasmid that recapitulates and completes all the properties of the pRS plasmids. The COSPLAY toolbox contains a basal collection of individual functional modules. Moreover we standardized a simple and rapid, software-assisted protocol which facilitates the addition of new personalized modules. Finally, our toolbox includes the possibility to select a genomic target location and to perform a single copy integration of the expression vector.

## Introduction

Over the last decade, the explosion of genome-wide analyses, as well as the rise of synthetic/systems biology has profoundly transformed the methodology used to study model organisms (Giaever and Nislow, 2014). Yeast has been a pioneering model in these emerging fields thanks to the versatility and the ease to perform genetic manipulations. High recombination rates, fast generation and crossing times are crucial features that allowed to create large collections of tagged and deleted strains (Howson et al., 2005; Huh et al., 2003; Winzeler et al., 1999) which proved to be very useful for systematic functional studies (Giaever et al., 2002; Hillenmeyer et al., 2008).

Whereas large strain collections are built using single-step PCR-based gene replacement techniques, the vast majority of classical genetic studies in yeast rely on the use of expression vectors, such as the YXplac (Gietz and Sugino, 1988) and pRS (Brachmann et al., 1998) plasmid series, which are based on pUC19 and pBluescript backbones, respectively. These vectors feature a number of unique cloning sites and commonly used auxotrophy selection cassettes (HIS3, LEU2, TRP1, URA3, ADE2 in the pRS system), which make them convenient for genetic manipulations of standard lab strains. There are three types of such plasmids with different replication properties: integrating (stable chromosomal integration), centromeric (episomal low-copy plasmid), and 2 microns (high copy episomal plasmid) (Brachmann et al, 1998).

Since the first release of the pRS series, several improvements were brought to these expression vectors: first, the development of antibiotic resistance markers, such as the KanMX module conferring resistance to Geneticin (G418) (Brachmann et al., 1998; Wach et al., 1994), expands its versatility of use as no auxotrophic mutation is required in the targeted strain. Second, a set of pRS-based plasmids containing standard controllable promoter sequences (GAL1, MET25) were constructed to standardize the means to ectopically drive the expression of a gene of interest in a controlled manner (Mumberg et al., 1994). This collection was later expanded to fine tune expression level in a synthetic biology context, by making the constitutive promoting sequences of the following genes available: TEF1, ADH1, TDH3, CYC1 (Buchler and Cross, 2009). Next, an updated version of the pRS series, called pRSII, was reported, which substantially increases the number of available selection markers (auxotrophic: HIS2 and ADE1; antibiotic: phleomycin, hygromycin B, nourseothricin, and bialaphos) (Chee and Haase, 2012).

Therefore, there are now a large number of potential combinations of plasmid features for ectopic gene expression. This ever growing number makes it unrealistic for research labs to maintain an up-to-date collection of plasmids. Yet, the complexity of today’s genetic studies, in which more than eight mutations or tags are routinely used within a strain (Lu and Cross, 2010), strongly pleads for a flexible expression system in which plasmid type (integrating, centromeric, etc..), selection marker, and gene promoters could be selected in a systematic and modular manner.

Since the late 1990s, several fast cloning strategies have been developed to ensure a quick and reliable assembly of DNA fragments within a destination vector. The first system released is known as the Gateway vector system (available from Life Technology), which uses a proprietary recombination scheme to assemble up to 4 inserts into a destination vector following a 3-step procedure (Hartley et al., 2000). Interestingly, specific Gateway destination vectors with widely used selection cassettes, tags and promoters have been built for budding yeast (commercially available on addgene.com) (Alberti et al., 2007). However, a major drawback is that Gateway is a patented system which is not open to modifications, and may involve expensive running costs. In addition, due to the large number of inserts available in yeast, it is still necessary to purchase a large collection of plasmids (288 plasmids) in order to avoid multiple sub-cloning steps, since combinatorial shuffling of inserts in one step is not possible.

As an alternative to Gateway, the BioBrick cloning format was developed as an open-source system intended to normalize plasmid construction for synthetic biology. It uses a DNA editing technique based on 4 restriction enzymes to iteratively assemble inserts in a standardized manner (Shetty et al., 2008). However, there are as many steps in the assembly process as the number of fragments to integrate in the ultimate destination vector. Therefore, swapping a single insert from a complex destination plasmid requires starting the assembly process from scratch.

To overcome these limitations, a novel strategy based on the exonuclease activity of polymerase (or using a dedicated exonuclease) was used to generate sticky ends in the destination vector as well as in the insert to be integrated. These methods, known as SLIC (Li and Elledge, 2007) or Gibson assembly (Gibson, 2009; Li and Elledge, 2007) provide a powerful way to assemble multiple DNA fragments within one destination vector in a single step, and have been followed by others that used a different implementation with a similar outcome (Quan and Tian, 2009; Zhang et al., 2012). Yet, in common to all these techniques, the assembly process relies on the annealing of several sequence-dependent single strands DNA pieces, thus making the assembly efficiency somewhat variable.

Another strategy developed recently is the Golden Gate system, which is based on the use of a type IIS restriction enzyme (such as BsaI) that cuts outside of its recognition sequence (Engler et al., 2008, 2009). Using an appropriate design of flanking sequences of the DNA fragments to be integrated, one can assemble a virtually unlimited number of modules in the right order within one reaction step. This technique has been further developed to allow large-scale construction of multigenic vectors using a multi-level assembly scheme (Sarrion-Perdigones et al., 2011; Weber et al., 2011).

The Golden Gate system has recently been adapted to yeast (Lee et al., 2015) with the generation of a collection of modules (i.e plasmid type, selection markers, promoters, etc…) that can be assembled in different combinations to produce expression vectors (up to 11 modules) as well as complex multi-gene assemblies for synthetic biology approaches. Here, we report the development of a simpler alternative plasmid architecture based on the Golden Gate system that allows both efficient assembly of individual modules into a destination expression vector and rapid generation of new custom modules using the MEGAWHOP technique. The resulting COSPLAY (COmbinatorial Swift PLasmid Assembly in Yeast) toolbox includes a collection of 26 modules that can be integrated into 6-module expression vectors that recapitulates and completes the properties of the pRS plasmid series. We have paid particular attention to make this toolbox straightforward to use and to expand it by developing custom Matlab software that automates the design of the tailed primers used to generate individual modules. In addition, unlike standard yeast integrating plasmids, our toolbox is designed to offer the possibility to ensure single copy integration of the expression vector to a target locus in the genome, independently of auxotrophic markers. The COSPLAY toolbox is available on Addgene as a package of 27 plasmids (URL link to be added) and the software can be download from github :https://github.com/gcharvin/cosplay.

## Results

### COSPLAY, a toolbox for rapid generation of yeast expression vectors

The object of the COSPLAY toolbox is to provide the possibility of easily, quickly and efficiently generate new expression vectors containing different combinations of functional elements or modules. Each of these modules can be selected from a collection of ready-to-use plasmids, or can be easily generated from cDNA, as described below. The organization of a yeast expression vector generated with the COSPLAY toolbox is based on six functional modules assembled in a specific order into a destination vector (pUC57; Figure 1A). Each position is committed to a specific role. The first (Figure 1) determines the mode of plasmid replication (i.e. integrating, centromeric, high copy (2 microns), or targeted to a specific locus in the genome). The second contains a promoter driving the expression of a cDNA located in the third position. If the cDNA lacks a stop codon, it can be fused to the subsequent module (Position 4), which could be either a fluorescent protein or a degron to decrease the stability of the resulting protein. The fifth position consist of a transcriptional terminator sequence followed by a selection marker (Position 6) used upon yeast transformation.

**Figure 1.**
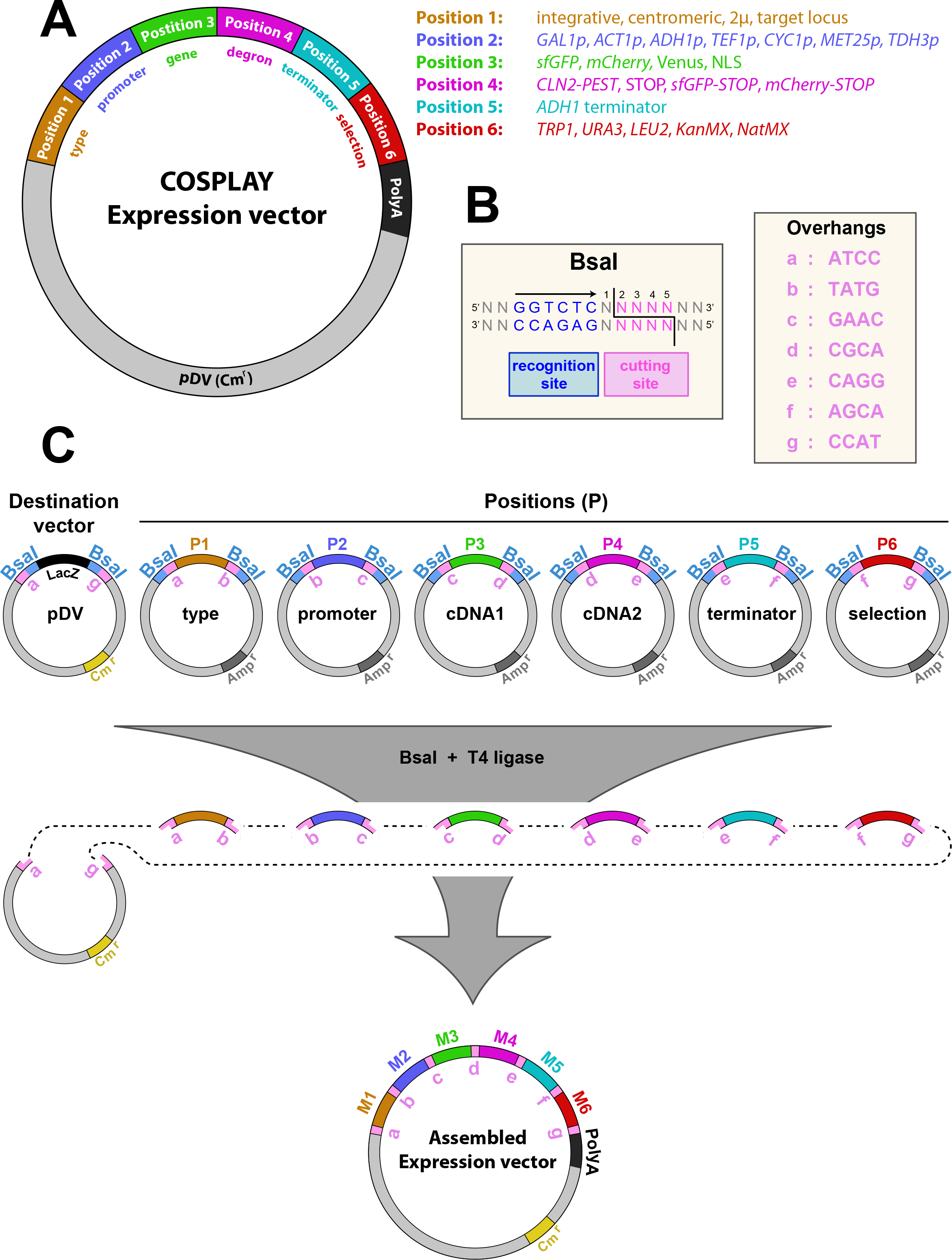
Plasmid architecture in the COSPLAY toolbox. A) Schematic representation of an expression vector assembled by COSPLAY composed of 6 independent modules. Each module represents a different type of sequence as indicated on the legend. B) Restriction site of the BsaI endonuclease and specific overhangs used; C) Principle of COSPLAY expression vector assembly. A destination vector (pDV) is mixed with 6 plasmids containing compatible module in a single tube. Expression vector is assembled through simultaneous digestion/ligation in the presence of BsaI and T4 DNA ligase. Specific overhangs that ensure that individual modules are assembled in the right order and orientation are indicated in pink.

The COSPLAY toolbox uses the Golden Gate cloning technology (Engler et al., 2008, 2009), which relies in the usage of Type IIS restriction enzymes (*i.e.* BsaI, BpiI, etc.). These enzymes cleave outside of their recognition site (Figure 1B), leaving a 4 bp 3’ overhang that can be manipulated to generate DNA ends (Figure 1B), which will dictate the compatibility between individual DNA fragments and the cloning order and orientation (Figure 1C). In addition, by placing the recognition sites at each extremity of the DNA fragment of interest (in opposing orientations), the restriction sites are lost upon ligation. Thus allowing to simultaneously carry out the restriction digestion and ligation reactions in a single tube (Figure 1C). The destination vector (pDV) has been engineered to contain a chloramphenicol resistance (CmR) and carries only 2 BsaI restriction sites (compatible with the module library) and flanking a LacZ cassette and located upstream of an SV40 polyA sequence (Figure 1C). This cassette is lost upon successful assembly of individual modules into the destination vector and allows performing a Blue/White screening to increase the cloning efficiency. The product of the restriction/ligation reaction is transformed into *E. coli* and grown on CmR plates. **Since the destination vector and the backbone vector for the module library contain different antibiotic selection markers (Table 1), one can easily select only for the destination vector only**. Individual white colonies are grown in liquid culture, plasmid DNA is prepared and vector integrity is verified by restriction digest using appropriate restriction enzymes and sent for sequencing. With the COSPLAY toolbox, any given construct can be assembled in 4 days with 100 % cloning efficiency (see cloning note on methods section).

### Combinatorial generation of a large number of expression vectors from a small collection of modules

We have generated a library of modules, cloned in specific positions that are readily available in the COSPLAY toolbox and which correspond to widely used sequences for yeast genetic engineering (Table 1).

Regarding position 1 (plasmid type), we have built modules that contain either a 2μ plasmid replication sequence (high copy number), a CEN-ARS sequence for transient extrachromosomal expression, or a nonfunctional short linker sequence to generate an integrating plasmid (in this case, homologous recombination leading to chromosomal integration is obtained by linearizing the plasmid within the selection marker). In position 2, we have generated widely used inducible or constitutive promoters, such as *GAL1p*, *MET25p*, *ACT1p*, *CYC1p*, *TEF1p*, *ADH1p and TDH3p*. Since classical expression vectors (pRS, pRSII,..) are often used to make fluorescent transcriptional reporters, we have generated modules in position 3 containing fluorescent proteins (i.e. superfolder GFP (sfGFP), mCherry, Venus). These sequences lack a stop codon and can thus be destabilized by adding a CLN2-pest degron cloned in position 4 or not, if a stop codon is cloned in position 4. Furthermore, to enable the fusion of a C-terminal fluorescent tag to a protein of interest, we have generated additional modules in position 4 encoding sfGFP and mCherry (bearing a STOP codon). We have generated a module in position 3 carrying a nuclear localization signal (NLS) sequence based on three repeated sequences from the SV40 virus, in order to allow the expression of nuclear fluorescent reporters (when cloned with sfGFP or mCherry in position 4). In position 5, we have cloned a single module which carries the *ADH1* transcriptional terminator. Finally, position 6 provides a large choice of universal yeast selection markers such as auxotrophy genes (i.e. *TRP1*, *URA3*, *HIS3*, *LEU2*) as well as drug resistance cassettes (i.e. *KanMX*, *NatMX*).

All modules (Figure 1A and Table1), in all positions, are compatible with each other within the framework of an expression vector containing 6 elements (Figure 1). Therefore, despite the relatively small collection of functional modules, the COSPLAY toolbox offers **extreme flexibility in the assembly of different expression vectors**.

### An efficient cloning method to generate new modules

In addition to the modules available in the COSPLAY library it is possible to easily generate new modules to meet user-specific needs (e.g. a specific cDNA, promoter, fluorescent protein, etc.). To this end, we have standardised a cloning-free two-step PCR process (Miyazaki, 2011) whereby the sequence of interest is amplified by PCR using appropriate primers containing bsaI sites with appropriate overhangs and 25 bp of the pUC57 multiple cloning site (Figure 2A). The PCR product of this first reaction is then used as a mega-primer to amplify the pUC57 destination vector, leading to the direct integration of the target sequence into pUC57 (Figure 2A). The pUC57 plasmid contains a *LacZ* cassette, which is rendered nonfunctional when a DNA sequence is cloned in its multiple cloning site, thus allowing Blue/White screening after transformation in *E. coli* (Figure 2C).

**Figure 2.**
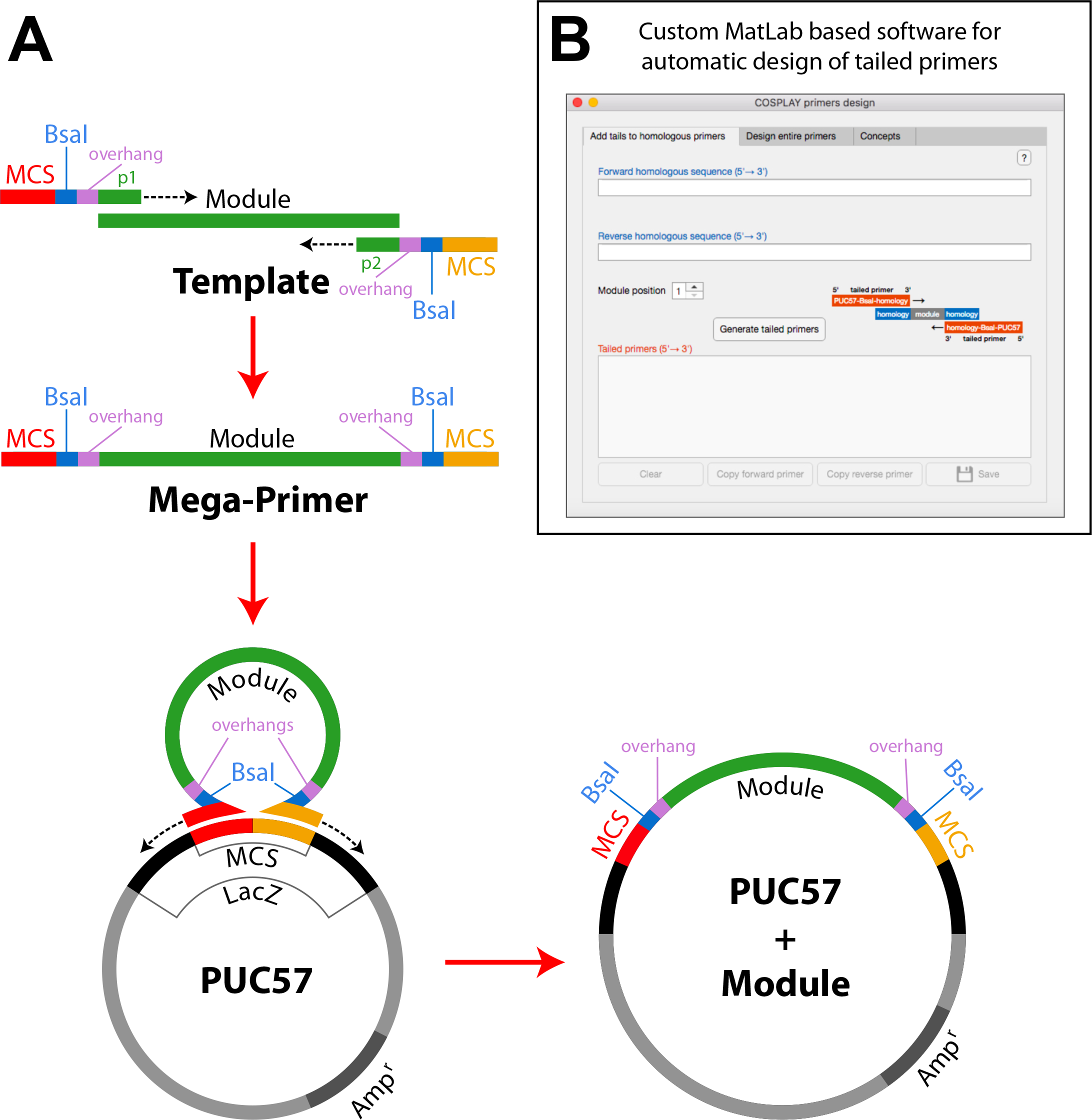
PCR-mediated generation of custom-made modules. A) Sketch showing the primer design and the two PCR steps required to clone a specific module (See supplementary Figure 2 for details on primer design); B) Snapshot of the Matlab graphical-user interface used to automate the design of primer tails required for the generation of module variants.

In order to facilitate primer design to generate new modules, we have developed a custom program with a graphical user interface in Matlab (Figure 2B) that provides assistance in this process: the user either enters the sequence of primers homologous to the sequence to be integrated in the module or the complete sequence of the module. For each module position (1 to 6), the program adds the primer tails and runs a battery of tests to evaluate the quality of the primers. In addition, the program identifies potentially conflicting BsaI restriction sites in the module sequence and proposes strategies to mutate them through fusion PCR. A comprehensive documentation of the program is provided as supplementary material, and the program can be downloaded from GitHub:https://github.com/gcharvin/cosplay.

### Generation of transcriptional reporters as an application of the COSPLAY toolbox

As a proof-of-principle we generated transcriptional reporter plasmids for the Gal1 and Cyc1 promoters using mCherry and sfGFP fluorescent proteins, respectively (Figure 3A and 3B). For the galactose-inducible promoter (Gal1p), cells were grown in SD medium supplemented with 2% dextrose, then transferred to a SD medium supplemented with 2% Raffinose and 1.5% Galactose before observation under the microscope for fluorescent protein expression. As expected, when the SV40 nuclear localization sequence was cloned in position 3, fluorescent signal was entirely nuclear (constitutively expressed with ACT1p-Figure 3C and 3D).

**Figure 3.**
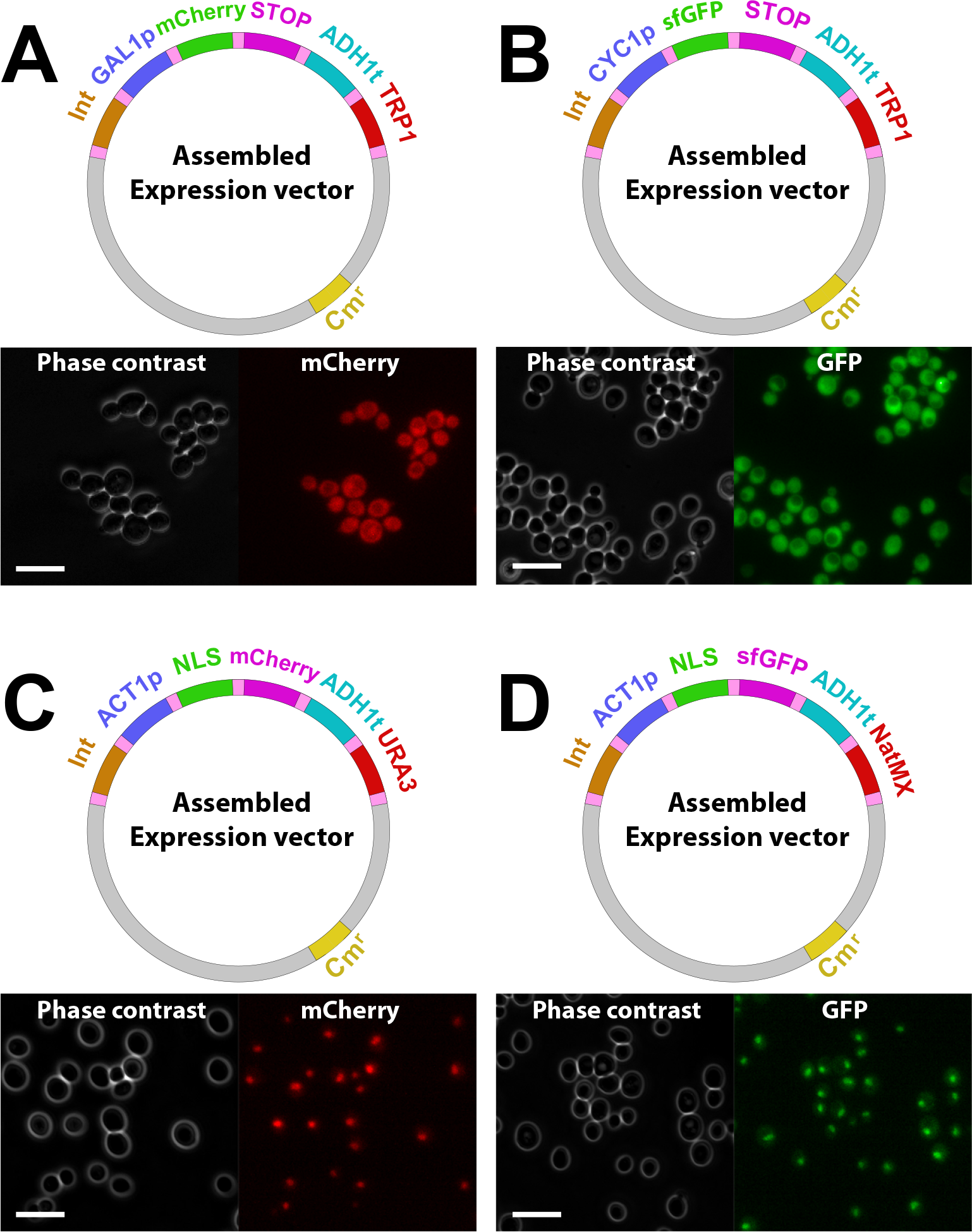
Proof-of-principle of the COSPLAY toolbox. Top panels: Maps of the different expression vectors generated by COSPLAY; Bottom panels: phase contrast and fluorescence images of cells transformed with the indicated plasmids.

### Single copy integration of plasmids using the COSPLAY toolbox

Quantitative studies in which ectopic gene expression is used require a precise control of plasmid copy number. Unlike centromeric and multicopy 2μ plasmid, integrative plasmids guarantee a stable expression over time. However, multiple integration events leading to variations in plasmid copy number across selected clones are not uncommon (Wosika et al., 2016). This is due to the fact that the integration site is replicated following the first homologous recombination, hence enabling subsequent integrations (Figure S1). Accordingly, it has been shown that the standard integrative plasmids pRS have a strong tendency to integrate multiple times at the same genomic locus while the single integration represents only 12.5% (when transformed with 1.5 μg of linearized plasmid) (Wosika et al. 2016). Therefore, the number of integrated clones must be assessed quantitatively either by directly scoring integration events (e.g. using southern blot) or by measuring the expression level of the gene of interest (e.g. using QPCR, western blot, cytometry, etc..).

To overcome this limitation, we developed a module in position 1 that ensures a single copy integration of the expression vector to a specific target region in the genome. This module is based on two inverted sequences (separated by the rare AscI and FseI restriction sites) which are homologous to a small genomic region (see Figure 4A and Methods). To facilitate module variant design, we have added an option in the software that takes this additional constraint into account for the generation of primers.

**Figure 4.**
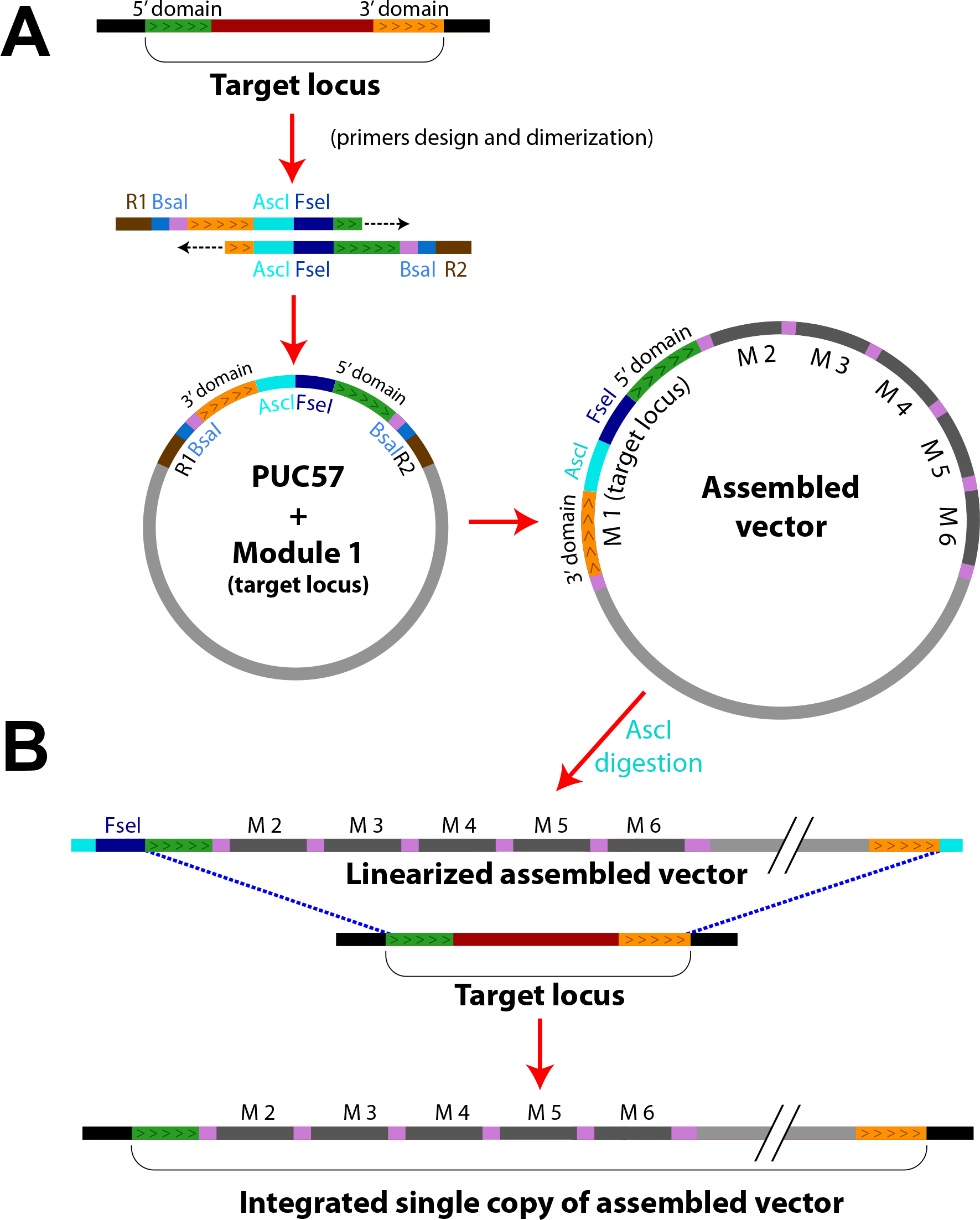
Principle of integration to a target chromosomal locus. A) A given locus is represented with its 5’ (green) and 3’ (yellow) domains. Primers are designed so that the two sequences are placed in reverse order (3’ region upstream of 5’), and AscI and Fse1 restriction sites are added in between these regions; B) The module carrying-plasmid and the final assembled vector are generated according to the standard procedure described in the text; C) Following AscI (or alternatively, FseI) digestion, the linearized plasmid is integrated at the target locus.

After AscI-driven linearization (or FseI if AscI is present in the inverted sequences), an assembled plasmid containing this module can be integrated at the homologous genomic locus of interest (Figure 4B). Subsequent plasmid integration is still possible, yet such event will only replace the already integrated plasmid copy (see Figure S1). Thus, only a single copy of a plasmid containing this particular modules should be inserted in the genome, unlike the chromosomal integration at the locus of an auxotrophic marker. Indeed, the fraction of single integration events has already been estimated to be close to 98% (Wosika et al,. 2016).

To check the validity of this approach, we have selected a target integration locus that fulfills the following criteria : 1) it must be far from telomeric and centromeric regions; 2) it must be in an ORF-free region of at least 1000 bp, flanked by ORFs that do not share the same regulation nor function. 3) It must be in a region of at least 200 bp devoid of protein binding sites and non coding RNA sequences. We found such loci in chromosome VI (position 260998 to 261148) and in chromosome IX (position 301997 to 302147). We have built the corresponding modules in position 1 using appropriate primer pairs as described above. Next, we have generated plasmids that contain these sequences, as well as a sfGFP transcriptional reporter of the Gal1 promoter and a Ura3 selection marker. These plasmids were linearized using either AscI (to target the specific loci on chromosome VI or IX) or EcoRV (to target the Ura3 locus), and all linearized plasmids were independently transformed in yeast (Figure 5A).

**Figure 5.**
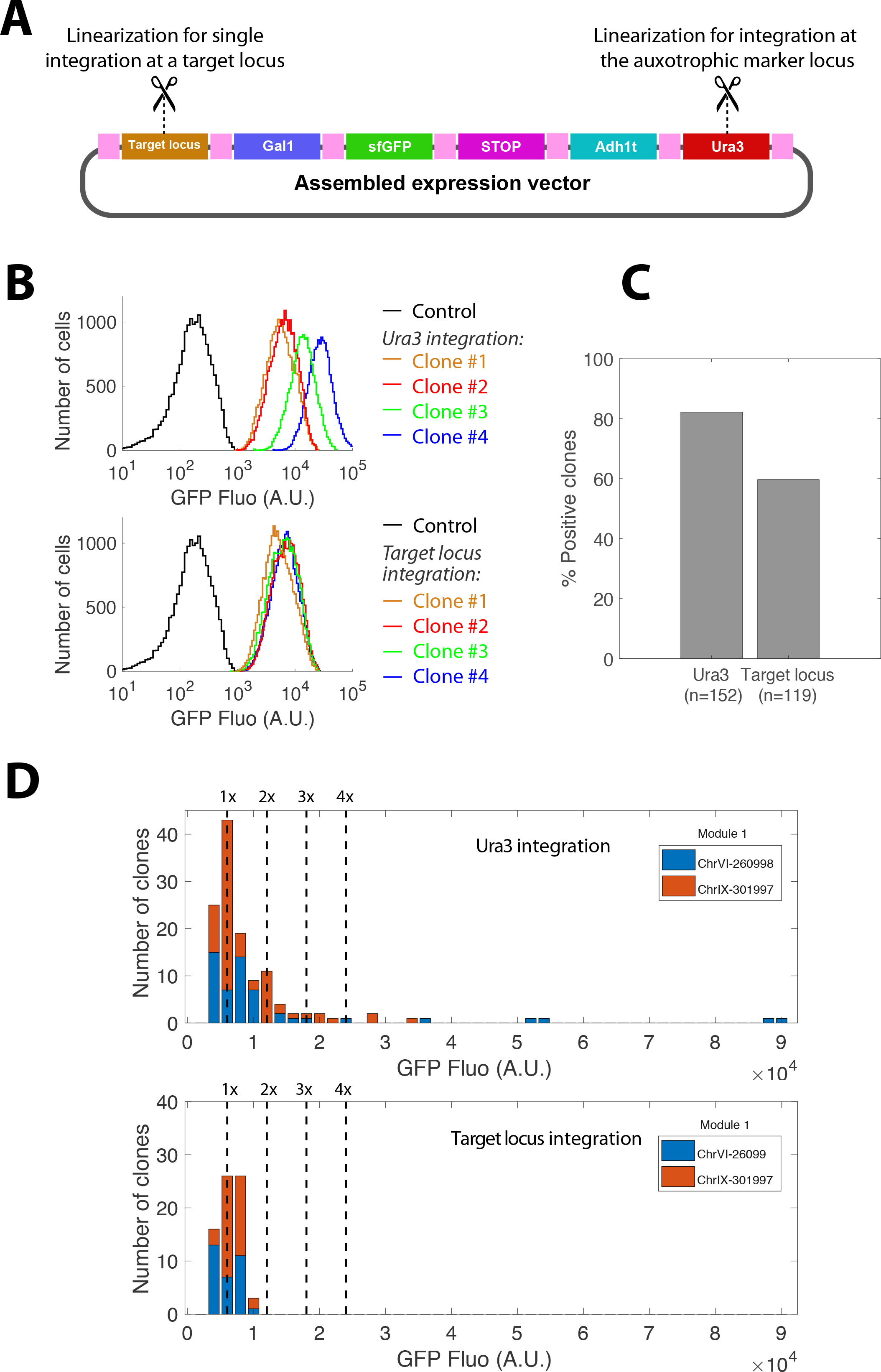
Single copy chromosomal integration. A) A sample plasmid containing the indicated modules is linearized by cutting either within the target locus (using AscI, see Figure 4 and corresponding text) or within the auxotrophic marker to compare the number of copy integrated at each locus. B) Flow cytometry analysis of GFP expression on yeast transformed with URA3, ChRVI or ChrIX-integrating plasmids. Each colored line represents the distribution of fluorescence obtained with a given clone. The black line represents the control. C) Fraction of positive clones (i.e. selected clones with a fluorescence higher than background) for strain transformed at the Ura3 versus a specific target locus integration; D) Distribution of median fluorescence of the positive clones (reported in C) selected after transformation at the Ura3 versus the specific target loci (ChrVI-260998 or ChrIX-301997); In the case of Ura3 (top plot), the histogram shows distinct peaks corresponding to 1- and 2-copy plasmid integrations as well as a higher number of integration; For the specific target loci (bottom plot), only the 1-copy peak is present.

To assess the number of plasmid copies respectively integrated at the chromosome VI, IX or at the Ura3 locus, we measured the protein expression level of more than 100 randomly selected clones of each type by flow cytometry (Figure 5B). As expected, recombinant clones exhibited a ~100-fold higher fluorescence level than the negative control (Figure 5B). Therefore, we quantified the fraction of recombinant clones for both Ura3 and integrations targeted to specific chromosomal loci (Figure 5C) and we calculated the median fluorescence level of each clone (Figure 5D). As expected, standard integration based on linearization within the Ura3 auxotrophic marker occasionally lead to multiple integrations (Figure 5D). In contrast, targeting specific loci on Chromosome VI or IX lead to single copy integrations (Figure 5D). Therefore, these results indicate that designing a custom module in position 1 targeting a specific chromosomal locus provides a straightforward way to ensure a single-copy integration.

## Discussion

In this study, we have developed a new toolbox called COSPLAY that provides fast and easy access to a high number of plasmid variants by handling the combinatorial part of the expression vector design. Our toolbox is based on a ready-to-use collection of individual functional modules and a one-step protocol to rapidly assemble a combination of modules into a functional expression vector. Each module can be assembled with any of the others modules leading to a large number of potential expression vectors.

A particular feature of the COSPLAY toolbox is that the collection of modules can be easily expanded using the MEGAWHOP protocol (Miyazaki, 2011). Here, specific sequences of interest (promoters, cDNAs, fluorescent reporters, epitope tags, etc.) are seamlessly cloned by PCR, thus producing the possibility to expand the flexibility of the COSPLAY toolbox at will. To streamline this process, we have developed a standardized protocol that, combined with our primers design software, makes the generation of new modules very straightforward. One limitation to keep in mind for module design is the hypothetical presence of BsaI restriction sites in the specific sequence of interest. In this case, the sites have to be muted and our software has been designed to assist users in this process.

While other combinatorial strategies for plasmid generation based on the Golden Gate system already exist for yeast (Lee et al,. 2015), we made the choice to develop a toolbox that is simpler, that yet recapitulates all the features of the pRSII series plasmids. Indeed, our toolbox is built on an expression vector scheme containing only 6 module types (versus 8-11 for alternative strategies). Moreover, we decided to use only one type IIS restriction enzyme (BsaI) for expression vector assembly. This further simplifies the toolbox since a smaller number of modules needs to be assembled, and there are less constraints regarding the presence of specific restriction sites in the module sequence.

The COSPLAY toolbox also allows for single integration using a single module that can be easily designed with our software, instead of the 3 modules required in alternative methods (Lee et al,. 2015).

To conclude, the COSPLAY toolbox is extremely flexible and efficient, simple of use, easily expandable and complementary to alternative Golden Gate cloning-based systems for yeast genetic editing.

## Material and methods

### Yeast strains and media

All strains were congenic to BY4742 (Winston et al., 1995; Brachmann et al., 1998) or MMY116-2C.

| Name | MA<br>T | Background | Genotype | Origin |
| --- | --- | --- | --- | --- |
| BY4742 | $\alpha$ | S288C | *his3Δ1; leu2Δ0; lys2Δ0; ura3Δ0* | Euroscarf |
| MMY116-2C | $\alpha$ | W303 | *leu2-3; trp1-1; ura3-1; his3-11; ADE2* | Mary<br>Miller |
| YAP43 | $\alpha$ | W303 | *leu2-3; trp1-1; ura3-1; his3-11 GAL1p-*<br>*mCherry::TRP1* | This study |
| YAP172-TRP9 | $\alpha$ | W303 | *leu2-3; trp1-1; ura3-1; his3-11*<br>*CYC1p-sfGFP::TRP1* | This study |
| YAP194 | $\alpha$ | S288C | *his3Δ1; leu2Δ0; lys2Δ0; ura3Δ0 ACT1p-*<br>*NLS-mCherry-URA3::targetVI;* | This study |
| YAP197 | $\alpha$ | S288C | *his3Δ1; leu2Δ0; lys2Δ0; ura3Δ0 ACT1p-*<br>*NLS-mCherry-NatMX::targetVI;* | This study |

The strains YAP194 and YAP197 were obtained by single integrations of reporter cassettes at a locus situated in chromosome VI (position 260998 to 261148).

Yeast cultures were grown in standard YPD medium (1% yeast extract, 2% peptone, 2% dextrose) or in minimal SD medium (0.67% Nitrogen base, 2% dextrose or galactose, amino acid mixture). Prototrophic transformants were selected for by plating on synthetic complete dropout plates (0.67% yeast nitrogen base, 2% dextrose, 2% agar) lacking the appropriate amino acid or nucleobase. Drug resistance transformants were selected on drug supplemented YPD plates.

To induce GAL1 promoter, cells were initially grown in SD medium containing 2% dextrose and then switched to SD medium containing 2% Raffinose and 1.5% Galactose.

### Module library construction

To generate individual modules we amplified the specific DNA sequence of interest by PCR using primers (Supplementary Figure 2) that contain 25 bp of the multiple cloning site (MCS) of pUC57 followed by the BsaI restriction site, a specific 4 bp overhang (Figure 1 and table below), which determines the cloning position. These sequence is in turn followed by 15-25 bp of the target sequence of interest (Supplementary Figure 2). PCR conditions: 98°C

| Position | Forward BsaI site | Reverse BsaI site |
| --- | --- | --- |
| 1 | 5'GGTCTC G **ATCC** 3' | 5'GGTCTC T **CATA** 3' |
| 2 | 5'GGTCTC G **TATG** 3' | 5'GGTCTC T **GTTC** 3' |
| 3 | 5'GGTCTC G **GAAC** 3' | 5'GGTCTC T **TGCG** 3' |
| 4 | 5'GGTCTC G **CGCA** 3' | 5'GGTCTC T **CCTG** 3' |
| 5 | 5'GGTCTC G **CAGG** 3' | 5'GGTCTC T **TGCT** 3' |
| 6 | 5'GGTCTC G **AGCA** 3' | 5'GGTCTC T **ATGG** 3' |

PCR products were amplified by PCR using the Q5 (NEB®) high-fidelity polymerase following standard conditions. The product of this first PCR was then used as a megaprimer to insert the target sequence into the pUC-57 destination vector through a second, PCR. To do this, 0.5 μl of the first PCR product are mixed with 50 ng of pUC-57 plasmid and a second PCR is carried out in a 25 μl reaction with the following conditions: 98°C (30 s) / **30 cycles - 98°C (10 s), 68°C (30 s), 72°C (30 s/kb)** / 72° C 2 min. The second PCR product is digested by the addition of 1 μl DpnI-FD (ThermoFisher®) for 1 hour at 37° C to destroy the methylated pUC-57 template plasmid. After digestion, 10 μl of this reaction are transformed into DH-5 α or TOP10 bacterial strains and plated on LB plates supplemented with Ampicillin and X-Gal for Blue/White screening. If repetitive sequences are to be cloned, the usage of the Stbl4 (Invitrogen) bacterial strain is recommended.

Note, for primer design it is mandatory to check whether the target sequence of interest contains BsaI sites. If so, these have to be mutated by PCR.

Note, all individual modules were tested functionally by assembling expression vectors followed by transformation in yeast (data not shown).

### Expression vector assembly

A combination of 6 modules was assembled into a destination vector (modified version of pUC-57 that does not contain BsaI sites and carries a Chloramphenicol resistance cassette instead of Ampicillin) by an all-in-one reaction of digestion/ligation.

The 6 selected plasmid modules are mixed together with the destination vector (pDV) and digested/ligated simultaneously in a 20 μl reaction. All plasmids should be at 40 fmol/μl. To prepare the corresponding plasmid dilution, use the following formula: [DNA in fmol/μl] = ([ng/μl DNA] x 1520)/Plasmid size (in bp). Reaction conditions: 1X NEB® Buffer 4, 1mM ATP, 20U BsaI-HF (NEB®), 400U T4 DNA ligase (NEB®) and H20 qsp 20μl. PCR program: 3 cycles - 37°C (10 min), 20° (10 min) / 20°C (50 min) / 50°C (20 min) / 80°C (20 min). 10 μl of this reaction are transformed into E. coli and plated on LB plates supplemented with Chloramphenicol and X-Gal for Blue/White screening.

### Yeast transformation

Yeast were transformed using methods involving lithium acetate, polyethylene glycol, denatured herring sperm DNA and sorbitol (Gietz and Schiestl, 2007; Gietz and Woods, 2001). Transformants were selected by spinning down yeast cells after a heat shock and resuspending them in sterile water before plating on the appropriate dropout or drug-selection medium. For drug selection, the yeast were resuspended in YPD and allowed to recover before plating.

### Microscope imaging

Cells were imaged using an inverted microscope (Zeiss Axio Observer Z1, or Nikon Tie). Wide-field epifluorescence illumination was achieved using an LED light source (precisExcite, CoolLed), and light was collected using a 100× N.A. 1.4 objective and an EM-CCD Luca-R camera (Andor).

### Fluorescence quantification by flow cytometry

Cells were grown in SD media until 0.5 OD and then analyzed by flow cytometry using FACS Celesta (BD Biosciences, San Jose, CA). Cytometry data analysis was performed using custom Matlab 2017b scripts.

### COSPLAY primers design software

The COSPLAY primers design software was developed in MatLab 2017b to simplify and to automate the addition of new custom modules to the COSPLAY modules collection. The software can be downloaded from github:https://github.com/gcharvin/cosplay (as a Matlab application).

The main functions of the COSPLAY software are:

1. Automated design of optimized primers for amplifying a specific target sequence. Several criteria (Primer Tm, length, self complementarity, 3’ complementarity, matrix complementarity, 3’ matrix complementarity, GC %, GC clamps, 3’ stability, 3’ GC%, primers pair Tm difference, cross complementarity, cross 3’ complementarity) are used for this optimisation. The basic rules of optimisation are similar to those of the Primer3 software (Rozen and Skaletsky, 2000).
2. Addition of pUC57 MCS sequences, BsaI sites and specific overhangs depending on cloning position. Detection of BsaI sites in the target sequence and replacement with silent substitutions whenever it is possible (or suggestion of alternative strategies to discard the sites).
3. Estimation of the best PCR temperature conditions. The Tm of the primers is calculated using an accurate thermodynamic approach (SantaLucia, 1998). This Tm is optimised for Q5 polymerase which is recommended to use in our protocol (if standard Taq polymerase is used, the Tm has to be lowered by 7°C). The effect of the buffer salt concentration is also calculated (Owczarzy et al., 2004). In case there are some nucleotides substitutions the mismatch effect on the Tm is also precisely estimated (Allawi and SantaLucia, 1997, 1998a, 1998b, 1998c, 1998d; Peyret et al., 1999).
4. Display of a full report about the quality of the primers pair when the entire tailed primers are automatically designed from a user-defined template or indication of potential quality problems when only tails are added to user-defined primers.

See additional COSPLAY software user guide as a supplementary text.

## Supporting information

User guide

## Supplementary Figure Legends

**Supplementary Figure 1.**
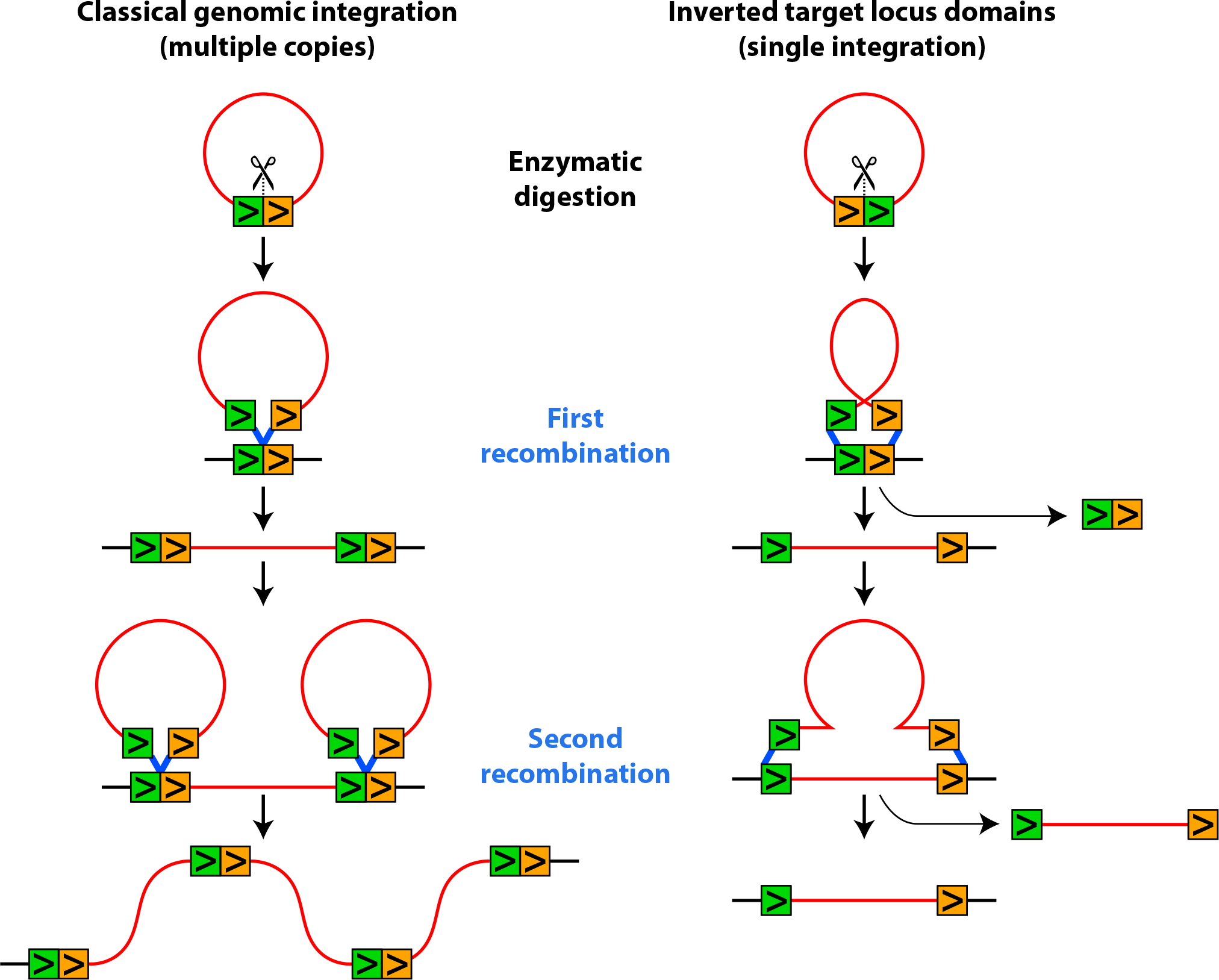
Principle of multi-copy versus single-copy chromosomal integration. Left: Chromosomal integration scheme after cutting the plasmid within a sequence homologous to a chromosomal region. Integration through homologous recombination duplicates the number of homologous regions that are available for further plasmid integration. Right: If homology regions are inverted on the plasmid, homologous recombination occurs in a way that preserve the number of homologous regions. In this case, further plasmid integration can only replace an existing insertion.

## Acknowledgments

We thank Maxime Diem and Vincent Hisler for contributions to this project during their stay in the lab as undergraduate interns. This work was supported by the ATIP-Avenir program (G.C.), and by grant ANR-10-LABX-0030-INRT, a French State fund managed by the Agence Nationale de la Recherche under the frame program Investissements d’Avenir ANR-10-IDEX-0002-02.

