## Supplementary material for "COSPLAY: An expandable toolbox for combinatorial and swift generation of expression plasmids in yeast": Table 1

COSPLAY library of modules:

| **ID** | **Position** | **Type** | **Description** | **Backbone** | **Resistance** |
| --- | --- | --- | --- | --- | --- |
| MV14 | 1 | plasmid type | *2µ* | PUC57 | Amp^R^ |
| MV8 | 1 | plasmid type | *flag (Yip)* | PUC57 | Amp^r^ |
| MV13 | 1 | plasmid type | *CEN* | PUC57 | Amp^r^ |
| MV46^2^ | 1 | plasmid type | *VI260998 (Yip)* | PUC57 | Amp^r^ |
| MV47^3^ | 1 | plasmid type | *IX301997 (Yip)* | PUC57 | Amp^r^ |
| MV1 | 2 | Promoter | *GAL1P* | PUC57 | Amp^r^ |
| MV2 | 2 | Promoter | *MET25p* | PUC57 | Amp^r^ |
| MV17 | 2 | Promoter | *ACT1p* | PUC57 | Amp^r^ |
| MV19 | 2 | Promoter | *ADH1p* | PUC57 | Amp^r^ |
| MV20 | 2 | Promoter | *CYC1p* | PUC57 | Amp^r^ |
| MV21* | 2 | Promoter | *TEF1p* | PUC57 | Amp^r^ |
| MV48 | 2 | Promoter | *TDH3p* | PUC57 | Amp^r^ |
| MV3^1^ | 3 | Reporter gene | *sfGFP* | PUC57 | Amp^r^ |
| MV4^1^ | 3 | Reporter gene | *mCherry* | PUC57 | Amp^r^ |
| MV22^1^ | 3 | Reporter gene | *Venus* | PUC57 | Amp^r^ |
| MV45^1^ | 3 | NLS seuqence | *aNLS3SV40* | PUC57 | Amp^r^ |
| MV5 | 4 | Cln2-PEST Degron | *CLN2-PEST* | PUC57 | Amp^r^ |
| MV23 | 4 | STOP codon | *STOP* | PUC57 | Amp^r^ |
| MV42 | 4 | Reporter gene | *mCherry-STOP* | PUC57 | Amp^r^ |
| MV44 | 4 | Reporter gene | *sfGFP-STOP* | PUC57 | Amp^r^ |
| MV6 | 5 | Terminator | *ADH1t* | PUC57 | Amp^r^ |
| MV7 | 6 | Selection marker | *TRP1* | PUC57 | Amp^r^ |
| MV10 | 6 | Selection marker | *URA3* | PUC57 | Amp^r^ |
| MV12 | 6 | Selection marker | *NatMX* | PUC57 | Amp^r^ |
| MV16 | 6 | Selection marker | *LEU2* | PUC57 | Amp^r^ |
| MV18 | 6 | Selection marker | *KanMX* | PUC57 | Amp^r^ |
| pDV | N/A | Destination Vector | *pUC57 with 2 BsaI sites flanking a LacZ cassette and carrying an SV40 polyA signal* | pUC57 | Cm^R^ |
| pUC57 | N/A | Module Cloning Vector | *pUC57 with no BsaI sites and carrying a LacZ cassette* | pUC57 | Amp^r^ |

^1^ MV3, MV4, MV22, MV45 modules do not have a STOP codon (fused to module 4)

^2^ MV46 corresponds to a target locus situated in chromosome VI (position 260998 to 261148)

^3^ MV47 corresponds to a target locus situated in chromosome IX (position 301997 to 302147)

* Note, plasmids using this module should be transformed into Stbl4 (invitrogen) or similar bacterial strains to avoid recombination between the Tet repeats.
