## Supplementary material for "COSPLAY: An expandable toolbox for combinatorial and swift generation of expression plasmids in yeast": User guide

When generating a new module the first step is to design primers that will amplify the module. These primers contain specific tails which drive the integration of the PCR product into a carrying plasmid and determine the future position of the module in a multi-modules assembly.The COSPLAY software propose 2 options to build primers: designing only the tails of the primers while the user freely choose the module homologous regions or designing the full tailed primers including optimized homologous regions.

In order to generate tailed primers when the homologous primer sequences are already available, the tab “Add tails to homologous primers” should be used:

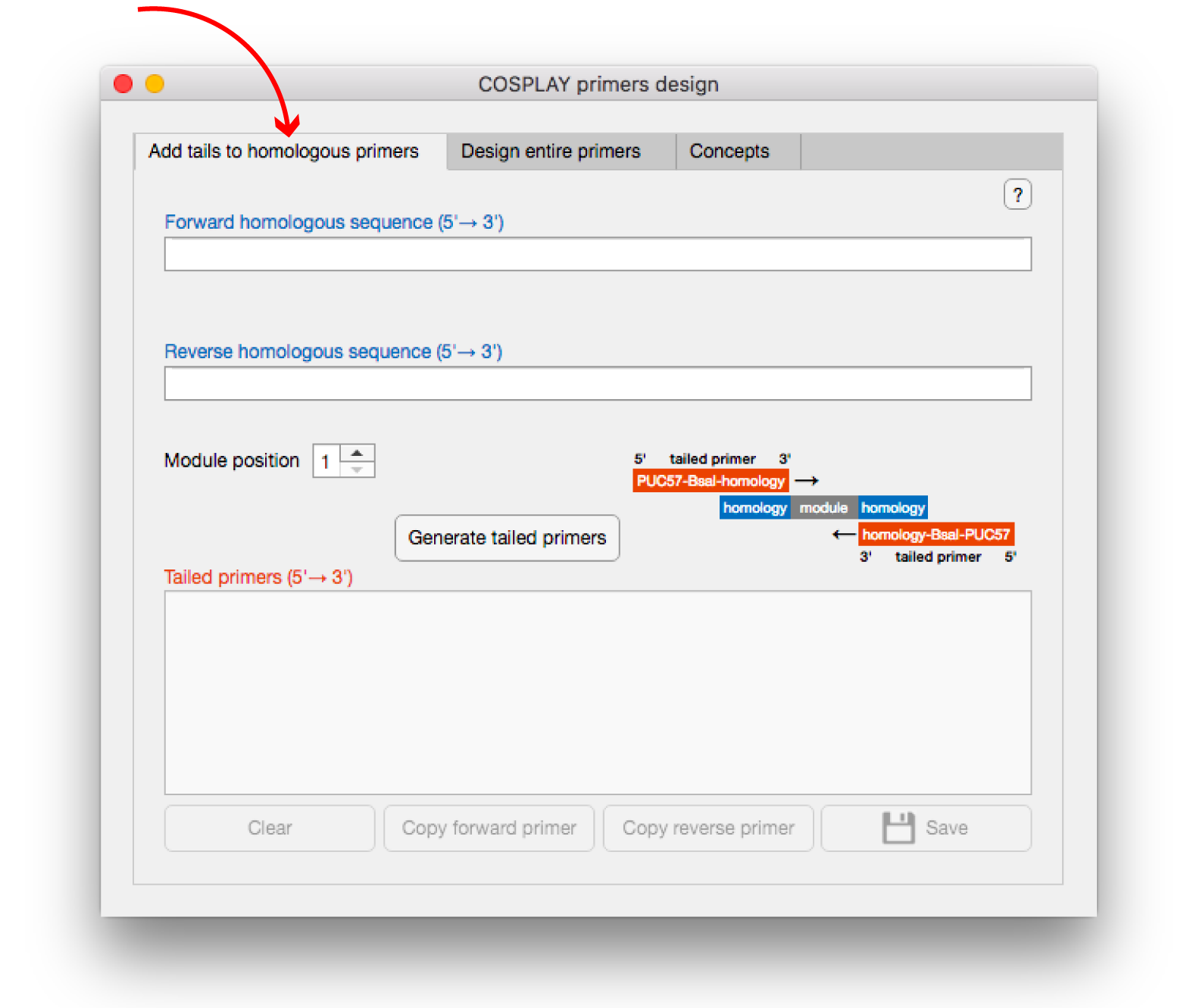

In this tab, the user is invited to enter 2 DNA sequences that are homologous to the 5’ and 3’ module regions. Only A/a T/t G/g C/c characters and blank spaces are allowed. A minimum length of 4 nt is required (minimum of 11 is recommended):

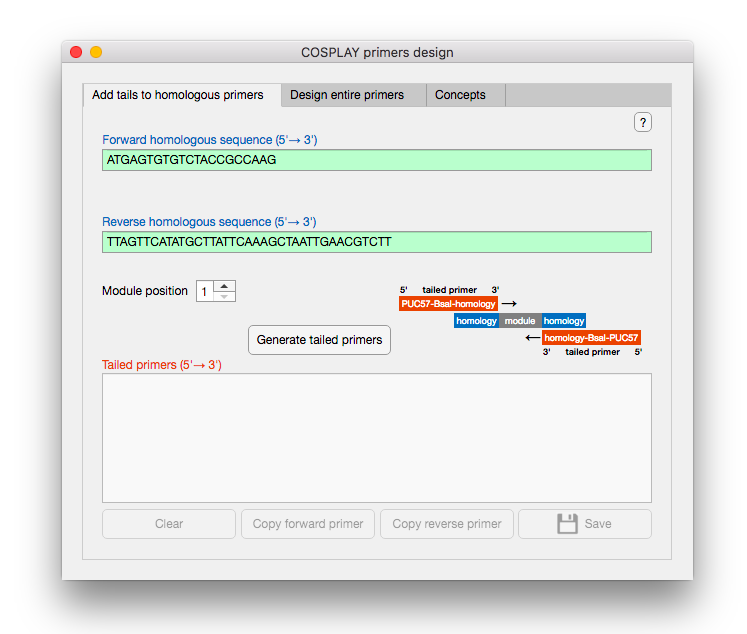

The next step is to choose what position the module should have in an assembled vector. In our COSPLAY toolbox the structure of an assembled vector is based on 6 different modules:

position 1 - plasmide type	position 3 - gene	position 4 - degron

position 2 - promoter	position 5 - terminator	position 6 - selection

The selection of the module position is achieved by using the corresponding spinner:

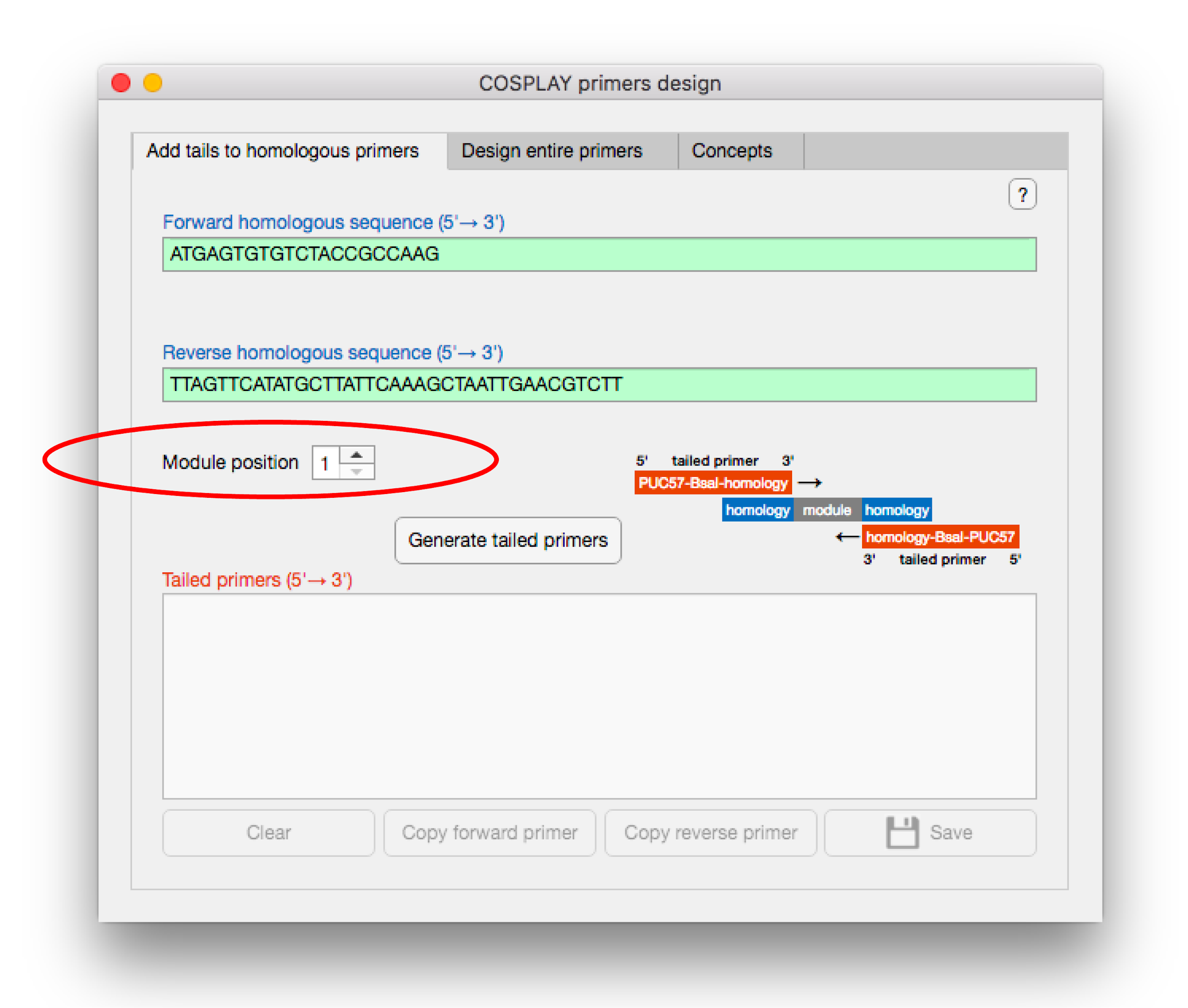

The button “Generate tailed primers” builds the tailed primers by integrating the provided information- homologous sequences and module position. The resulting sequences are displayed in the text area on the bottom. The primer Tm for Q5 pol is also shown:

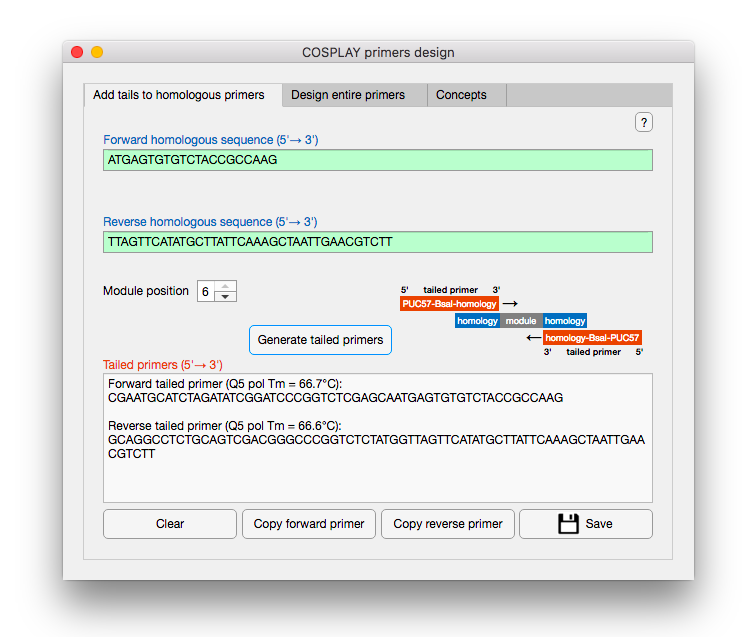

The sequences of the tailed primers generated by the COSPLAY software can be directly copied in the clipboard or saved as text file by using the corresponding buttons under the results text area.

Additional option is available for module position 3. Indeed, the module 3 may contain a gene that has to be fused to a destabilizing sequence contained in module 4. Therefore, the linker site between module 3 and 4 should be adjusted to preserve the reading frame. If no fusion is needed the standard linker sequence is [module 3]-^5’^CGCA^3’^-[module 4]. In the case of fusion the linker is extended by a gly-gly bond: [module 3]-^5’^GGAGGCGCA^3’^-[module 4]. To indicate that fusion is required the checkbox “Fused to module 4” should be checked:

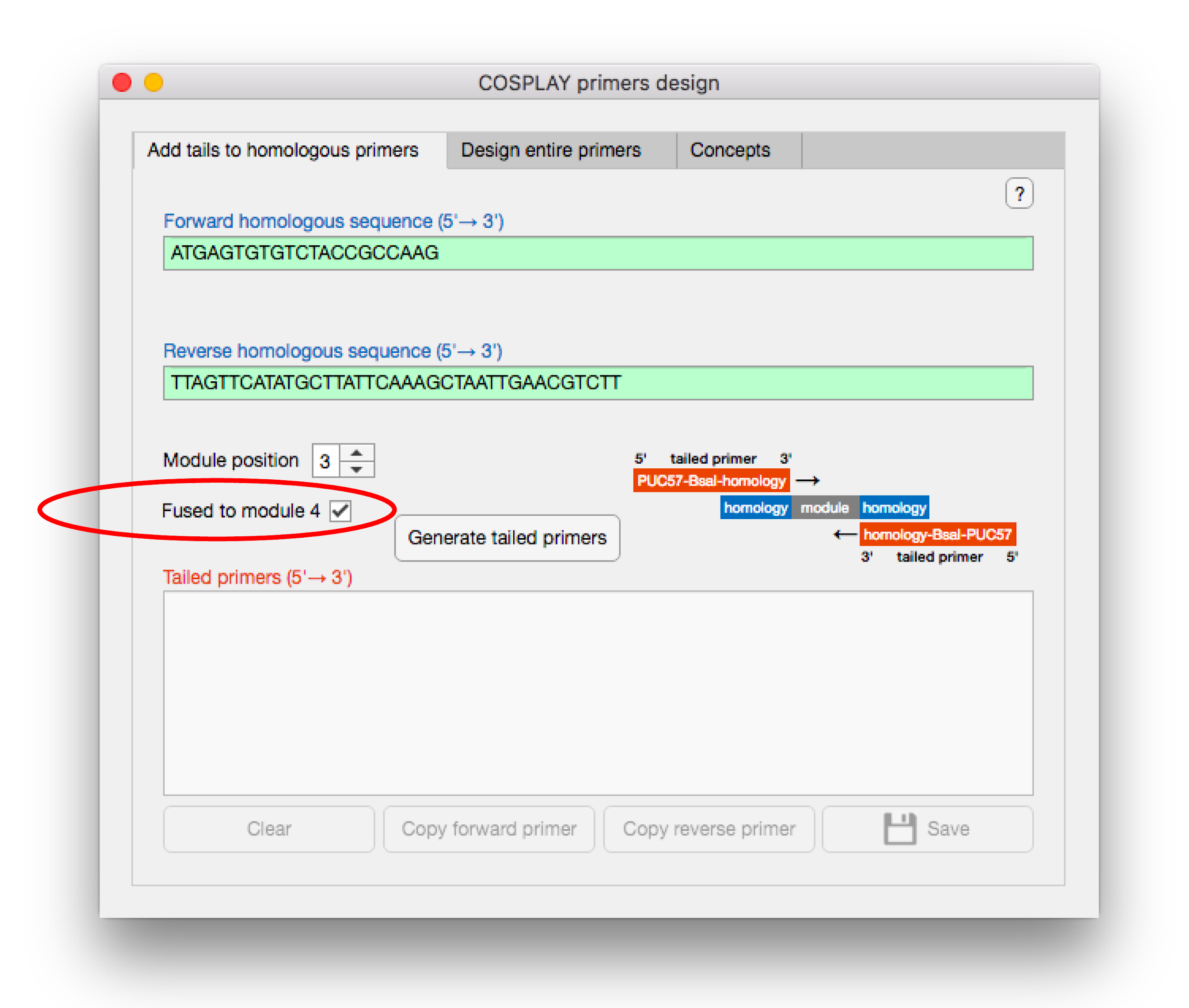

If the fusion option is validated the software tests whether a STOP codon is present in the reverse primer and eventually propose to eliminate it:

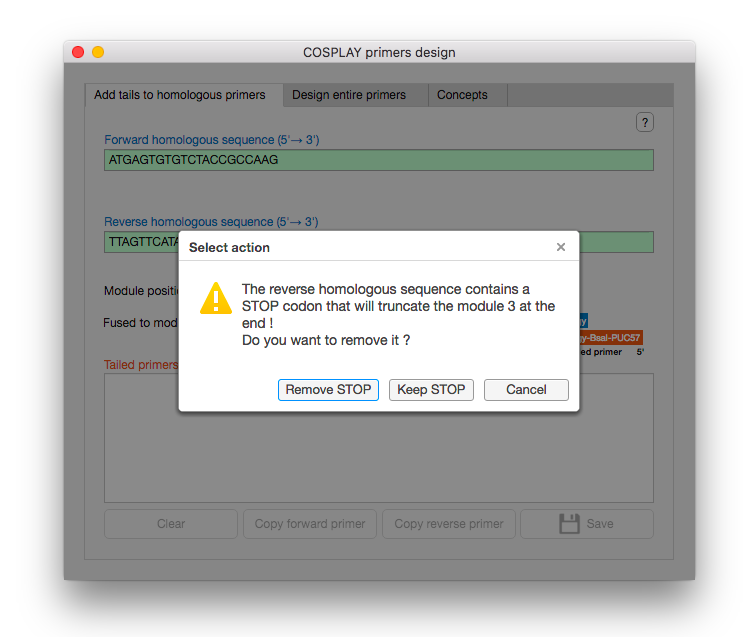

One limitation of the COSPLAY toolbox is the presence of additional BsaI sites in the modules sequence. Such sites have to be muted to avoid unintended cutting during assembly reactions (BsaI digestion + T4 ligase activity). Our software automatically detect the problematic BsaI sites present in the tailed primers and propose several silent substitutions depending on the reading frame:

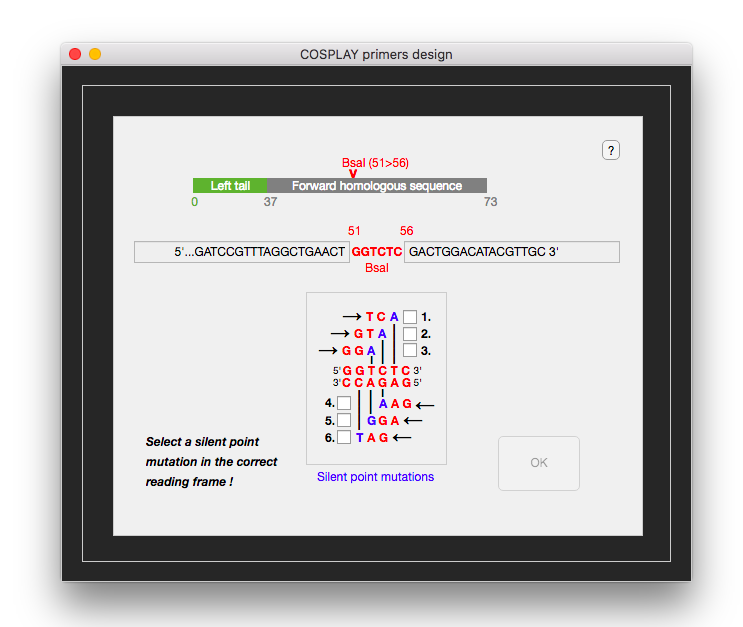

If the user can not provide the homologous sequences the tab “Design entire primers” may be used to find optimized homologous primers that amplify a module:

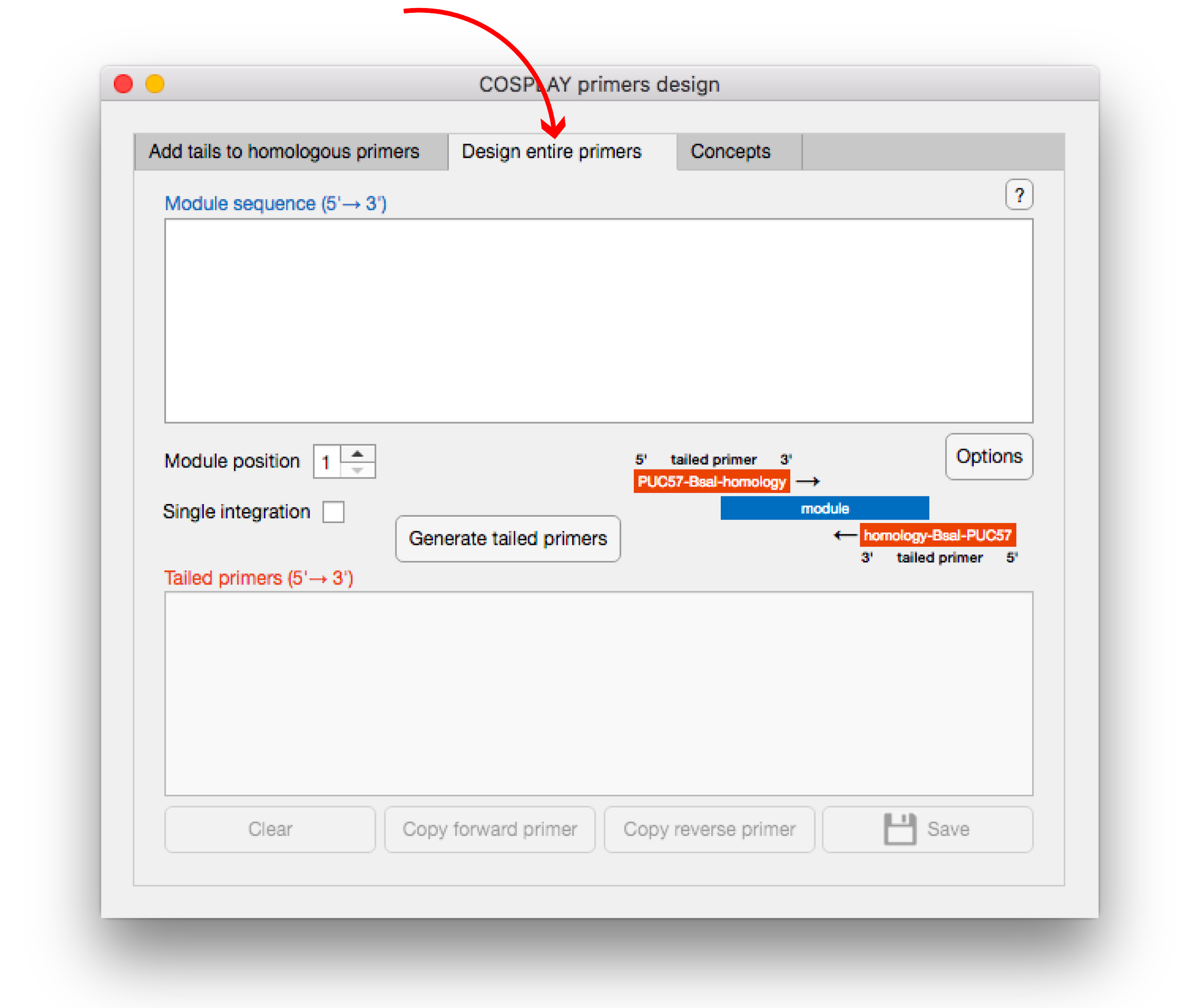

In this tab, the module sequence has to be communicated in order to generate the primers. For this the upper text area should be used:

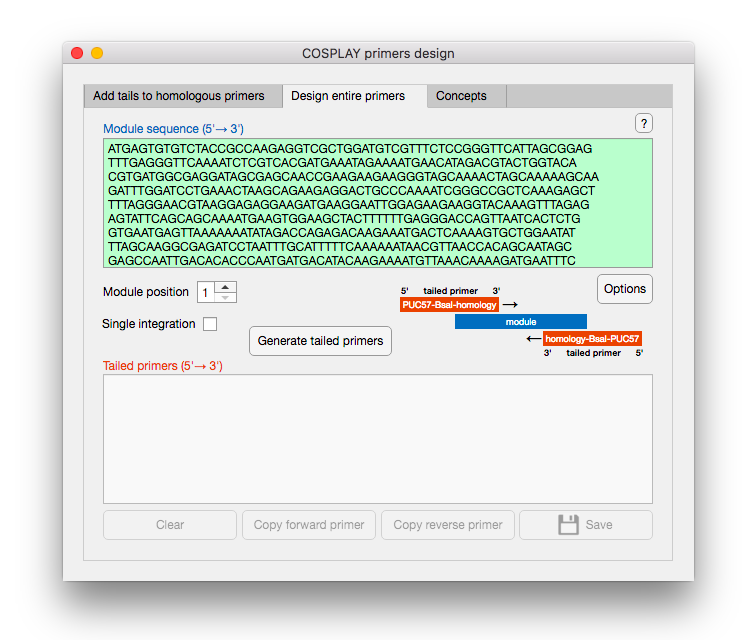

The forward and reverse homologous primer sequences always start respectively at the first (5’ end) and last (3’ end) nucleotide of the module. This guarantees that the entire module sequence will be amplified. The length of the homologous sequences is optimized to obtain the most efficient primers pair. Many criteria are considered during the optimisation process. The balance between different criteria is ensured via a system of weights that determine the importance of each criterion. The weights are numbers that multiply the optimisation effect of the criteria. Default values for all weights is 1 but this can be modified [0;+∞] by using the button “Option” situated under the module sequence text area. This button opens a new window that allow to change the weights as well as the optimal Tm:

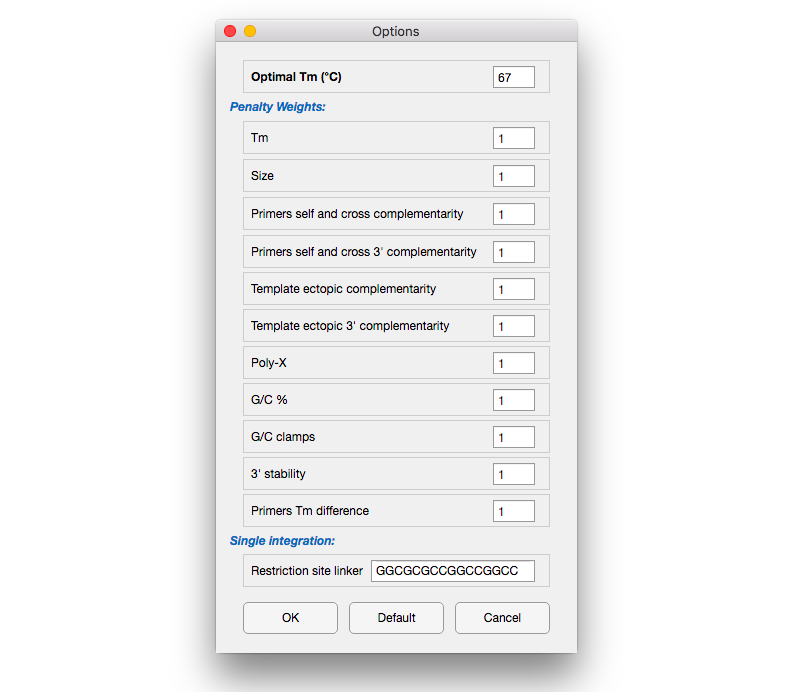

After the module sequence is communicated the user has to select the module position which determine the sequences of the tails that will be added to the homologous sequences in order to generate tailed primers. The button “Generate tailed primers” integrate all the input information to build the entire primers. The sequences of these primers and corresponding Tms are displayed in the bottom text area. In addition a full report on the primers quality is showed. In this report the different optimisation criteria are evaluated and results are based on 3 categories- “Ok”, “Mediocre” and “Bad” :

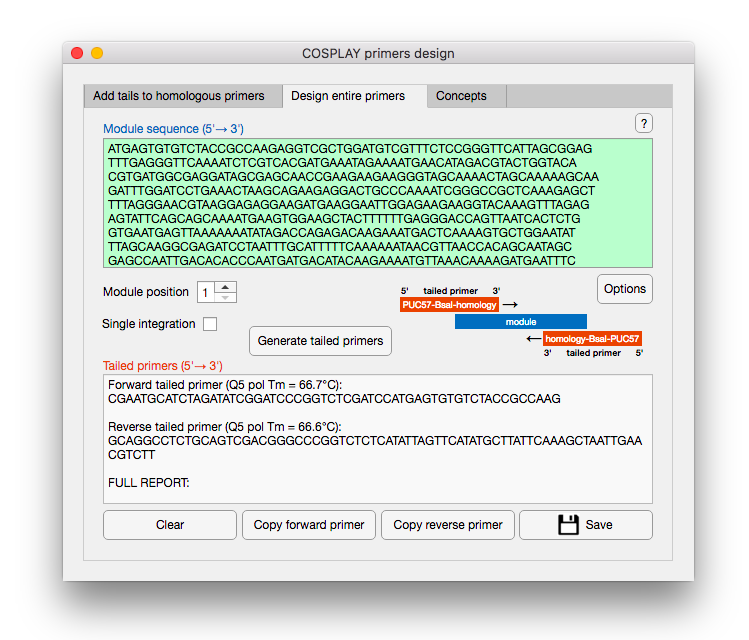

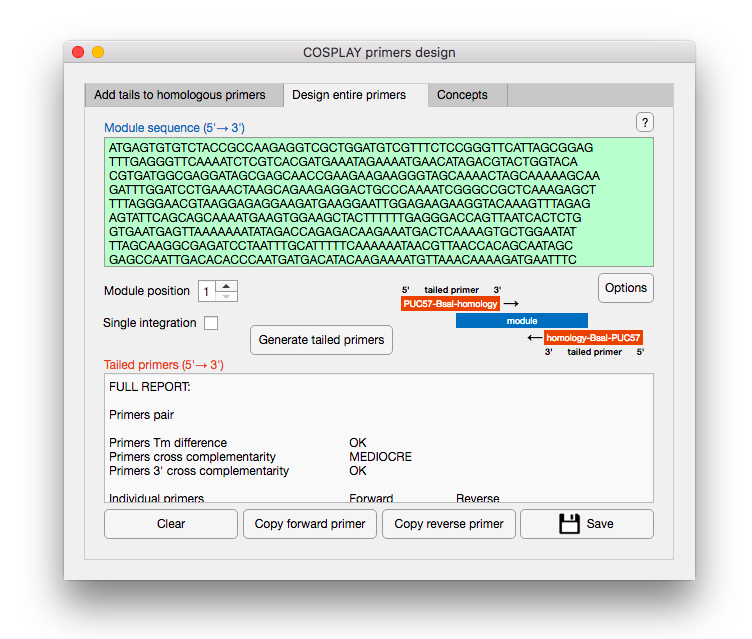

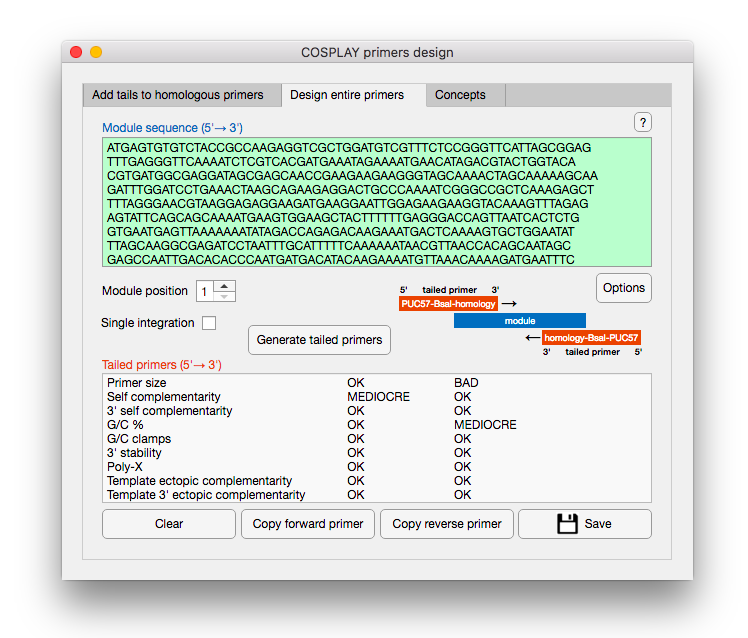

The forward and reverse tailed primers can be copied or exported as a text file:

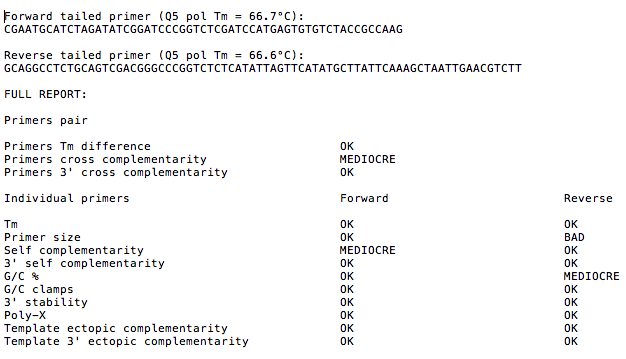

When the selected module position is 1 there is an additional checkbox determining whether the module is intended to perform single integrations.

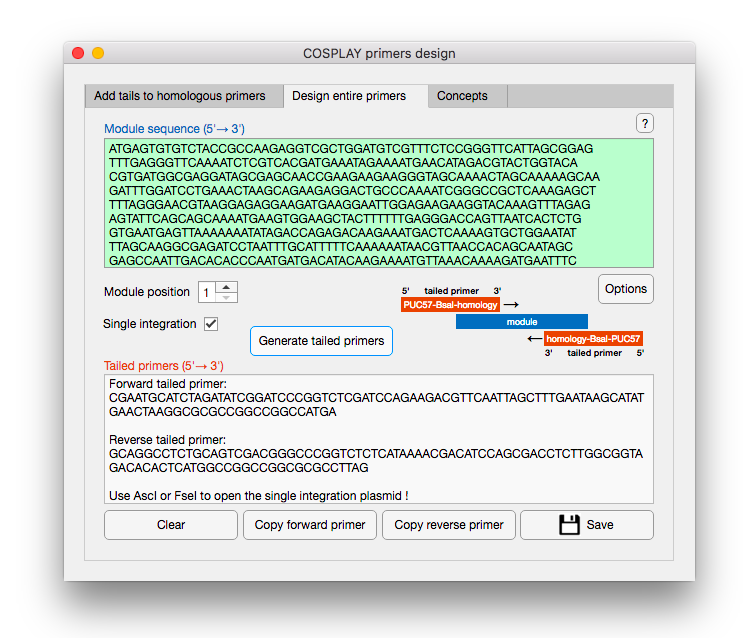

Indeed, the COSPLAY toolbox provides the possibility to do single genomic integrations of assembled vectors through a specific module 1. This module is based on 2 sequences homologous to the 5’ and 3’ regions of a target locus. The particularity ensuring the single integration is that these sequences are inverted (the 3’ region is upstream the 5’ region). If the checkbox “Single integration” is validated the software will generate the primers required for the production of a module 1 containing inverted 5’ and 3’ target locus regions (40bp each). The sequences are separated by a cassette containing AscI and FseI restriction sites used to open the plasmid prior to the genomic integration. These primers share 20bp of homology and should be directly dimerised by PCR (matrix not required!):

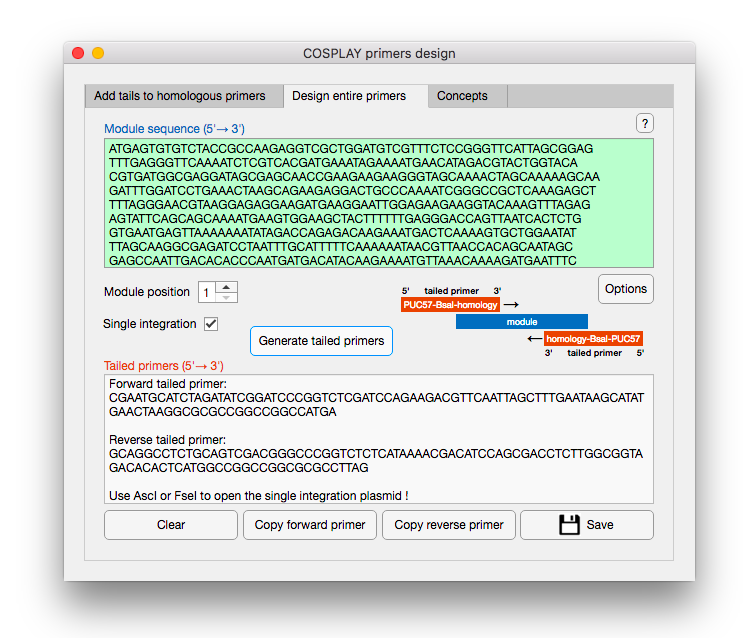

In case AscI and FseI are both present in the module 1 sequence outside the restriction cassette or in other modules that will be assembled with the module 1 it is possible to modify the restriction sites of the cassette through the button “Option”:

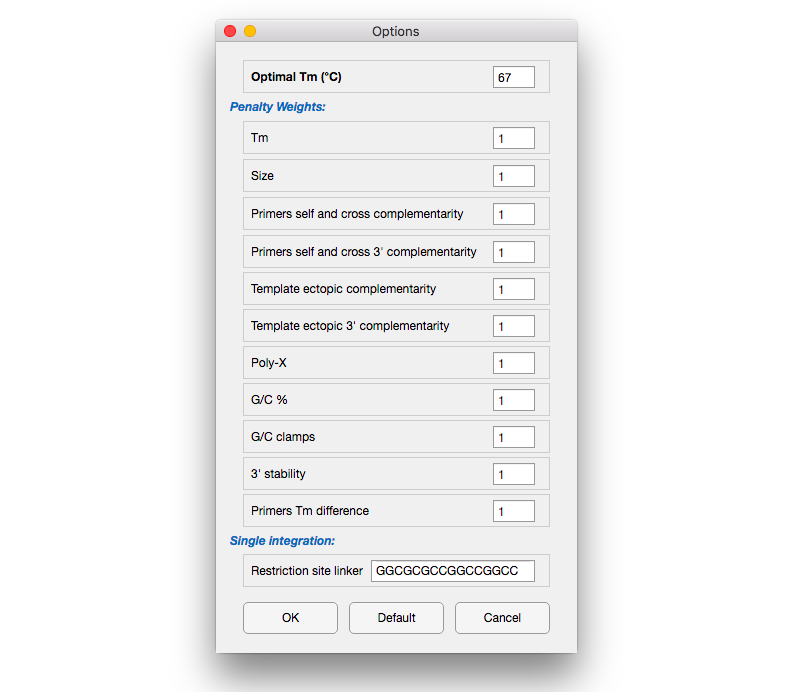

The last pad of the software contain informations about the main principles of the COSPLAY tool box:

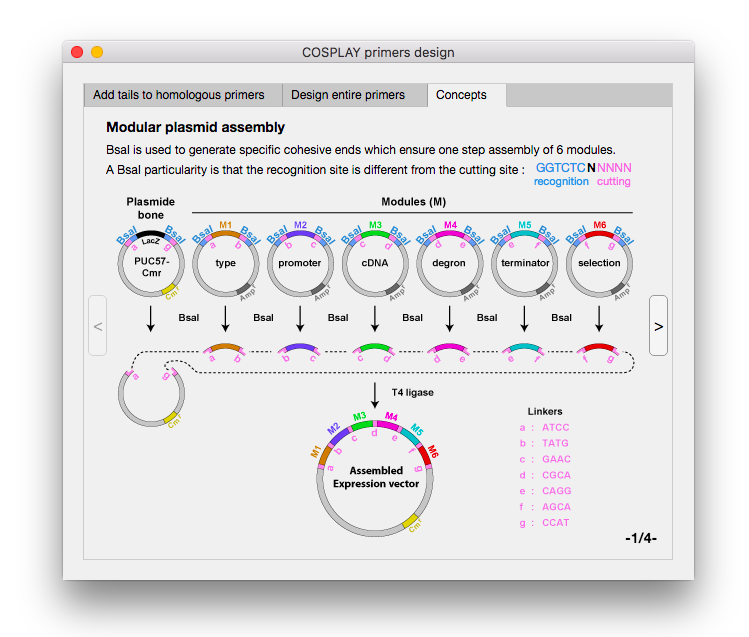

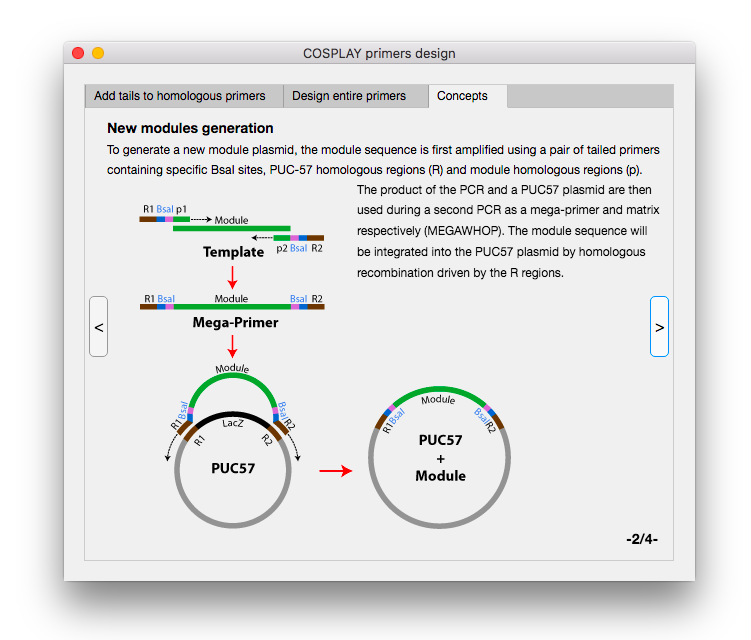

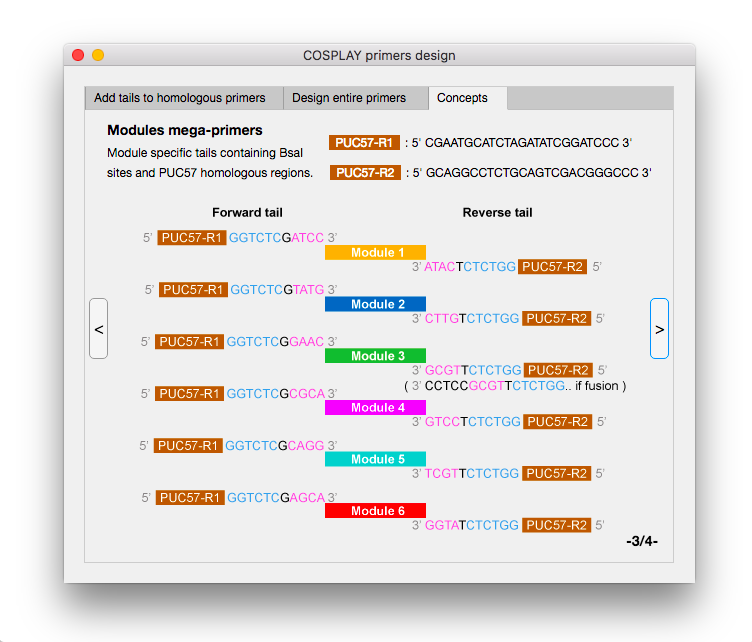

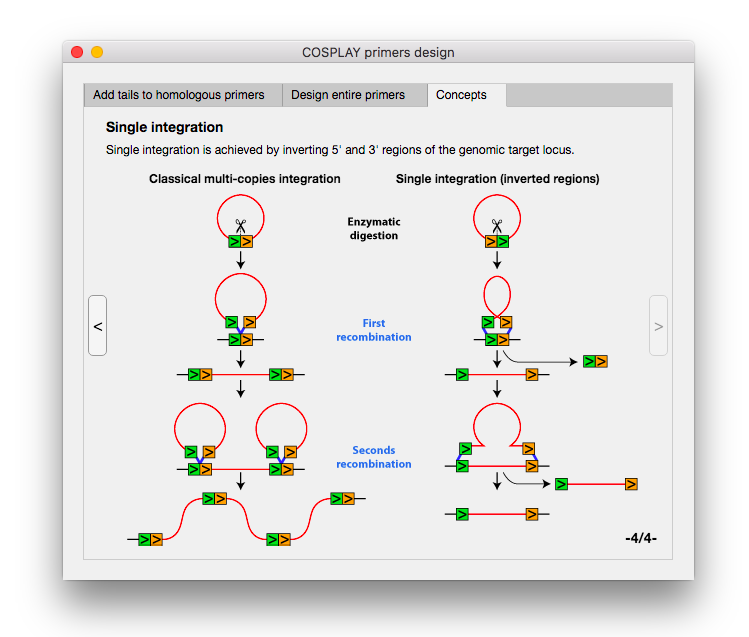
